# Alternative Polyadenylation alters UBASH3B-ZNF652 competition to involve in Triple-Negative Breast Cancer

**DOI:** 10.1101/2022.12.27.522044

**Authors:** Sijia Wu, Changbao Hu, Xiaoming Wu, Xiaobo Zhou, Liyu Huang

## Abstract

Alternative polyadenylation (APA) is an RNA-processing mechanism which may affect gene expression. Its involvements in cancers including the triple-negative breast cancer (TNBC) have been demonstrated in previous studies. Given the lack of biomarkers for TNBC, in this study, we were committed to finding novel biomarkers related to TNBC prognosis from the perspective of APA-mediated microRNA regulation. For this goal, raw bulk RNA sequencing data was collected for the identification of nine survival-related APA events by multivariate Cox regression analysis and forward-selected Likelihood Ratio Test. They showed good prognosis ability on TNBC risk. Of them, the APA event in *UBASH3B* was proposed as a novel potential biomarker. This event disturbed microRNA regulation on its host gene and other competing tumor genes to possibly involve in the pathogenesis of TNBC.

## INTRODUCTION

Triple-negative breast cancer (TNBC) is a subtype of breast cancer that lacks the expression of the estrogen receptor (ER), progesterone receptor (PR), and *HER2* ([1]). Due to its more aggressive behavior, poorer prognosis, and lacking of targeted therapies, TNBCs treated by chemotherapy are associated with a high rate of relapse ([2, 3]). For more precise and efficient therapies, previous researchers proposed several biomarkers. For example, tumor protein 53 gene (*TP53*) targeted compounds could restore its wild-type attributes to suppress tumor growth in breast cancer cells ([4]). Another gene of *VEGF* can mediate the signaling of angiogenesis associated with TNBC treatment in several studies ([5–7]). And the positive AR patients may have a more favorable prognosis for TNBC ([8–11]). These biomarkers may be insufficient and evidence-conflict ([12]). Thus, a novel kind of biomarker is still required for personalized treatment.

Alternative polyadenylation (APA) is a major mechanism of gene regulation employed frequently in almost all eukaryotes ([13]). As a main component of APA events, the APAs in 3’-UTRs can affect transcript abundance, control cellular localization, and interfere in microRNA (miRNA) regulation ([13]). Its roles in multiple tumor-related biological processes have been demonstrated in previous studies. Especially, shorten 3’-UTR of *CCND1* caused by APA events can promote the proliferation of breast cancer cell ([14]). And the cleavage factor Im 25 (CFIm25), a key factor for the selection of poly (A) sites, can affect oncogenes expression levels and tumor expansion in breast cancer ([15]). Given the potential of APA in cancers, in this study, we aimed to identify functional APA events associated with TNBC survival.

To realize this, we did as follows. First, survival APA events were found by univariate Cox Regression Analysis and forward-selection Likelihood Ratio Test (LRT) sequentially. Second, these survival APA events were used to build a risk signature. Its performance in TNBC prognosis was evaluated by KM survival and ROC curves. Next, the relationships of APA events with miRNA binding and gene expression levels were analyzed together to reveal the possible mechanisms of APA events in TNBC. Finally, an APA biomarker was popped up as the potential therapeutic target.

## MATERIALS AND METHODS

### Samples Collection

There were 352 TNBC patients from Fudan ([16]) and 108 TNBC patients from TCGA database that were included in this study, as shown in Table 1. Available data included their raw bulk RNA sequencing data for gene expression quantification and APA detection. Additionally, there were also clinical information such as age, Relapse-free survival (RFS) status, RFS time, tumor (T) stage, and node (N) stage for the clinical relevance of APA events.

**Table 1.** Clinicopathological characteristics of TNBC patients.

| DATASET | Fudan |  | TCGA |  |
| --- | --- | --- | --- | --- |
| Characteristics | detail | cox: HR(*P*) | detail | cox: HR(*P*) |
| **Age** |  |  |  |  |
| mean | 53.38 | 0.99(0.376) | 54.53 | 0.98(0.4055) |
| median | 54 |  | 52.5 |  |
| variance | 128.57 |  | 149.25 |  |
| **T stage** |  |  |  |  |
| 1 | 128(36.47%) | 1.42(0.228) | 26(24.07%) | 1.61(0.1195) |
| 2 | 215(61.25%) |  | 68(62.96%) |  |
| 3 | 8(2.28%) |  | 10(9.26%) |  |
| 4 | 0(0%) |  | 4(3.70%) |  |
| **N stage** |  |  |  |  |
| 0 | 209(59.71%) | 2.01(<0.01) | 67(62.04%) | 1.64(0.0976) |
| 1 | 91(26%) |  | 25(23.15%) |  |
| 2 | 33(9.43%) |  | 12(11.11%) |  |
| 3 | 17(4.86%) |  | 4(3.70%) |  |

### Detection of Candidate APA Events

To detect APA events, Quantification of APA (QAPA) ([17]) together with Salmon ([18]) were applied to calculate proximal poly(A) usage (PPAU). Moreover, to estimate the correlations between APA and gene expression, we used Spliced Transcripts Alignment to a Reference (STAR) ([19]) and RNA-Seq by Expectation Maximization (RSEM) ([20]) to quantify transcript abundances. During the process, GRCh37 was used as the reference genome.

Based on the APA events of samples from TCGA and Fudan, we checked their correlations. As shown in Extended Data Figure 1, the consistency of these two datasets revealed the co-pathogenesis of TNBC in different cohorts. Further, the APA events with mean PPAUs larger than 0.05 and standard deviation bigger than 0.01 were selected as candidate APA events for the following analysis.

### Identification of Survival related APA Events

To select APA events associated with survival, we first defined informative APA events by random sampling and univariate COX regression analysis (*P* ≤ 0.1) with the R package of “clusterProfiler” ([21]). The detailed process was described in the Algorithm in Supplementary File. To further reveal the associations between APA and breast cancer, we collected breast cancer related genes from DisGeNET ([22]), ONGene ([23]), OMIM ([24]), TSGene ([25]), dbVar ([26]), and ClinVar ([27]) databases.

Based on these informative APA events, we employed a forward-selected LRT to further identify survival APA events. It was described in the subsequent sections of Supplementary File Algorithm. Specifically, we first sorted the informative APA events in descending order according to the occurrences of APA events in the random sampling experiment. Then, the top 5% informative events were sequentially incorporated into the cox proportional hazards model (“rms” package) ([28]). During the incorporation process, LRT was used to determine whether the current new APA event was retained as a potential survival biomarker.

### Generation and Evaluation of Risk-Signature

With the survival APA events, a regression model ([29]) was used to generate a risk signature based on Fudan dataset as follows. First, multivariate COX regression analysis was performed on these survival APA events to estimate their associations with survival. Their contributions and PPAU values were combined for a risk signature which was normalized by Z-score standardization method. This risk signature was then used to classify patients with high and low risks using the optimal threshold determined by X-tile approach ([30]) based on Fudan dataset. To verify the robustness of this model, parameters and threshold value used in Fudan dataset were directly applied in TCGA dataset. The effectiveness of this risk signature in RFS prognosis was then evaluated by KM survival curves to classify the survival differences between high and low risk groups across different clinical subgroups.

### Exploration of the APA Pathways associated with TNBC Prognosis

The usage of proximal APA sites may lose potential miRNA regulation on their host genes ([13, 31, 32]). To investigate this, TargetScan ([33]) was applied to detect the loss of miRNA binding targets caused by proximal APA events. The changes of miRNA regulation were confirmed by the correlations between the PPAU values of the survival related APA events and expression levels of their host genes. Besides, to discover the further effects of the survival related APA events on other genes, we also analyzed the relationships between these APA events and the genes that compete with their host genes. Last, we performed enrichment analysis on these genes to reveal the probably involved biological functions of these APA events in TNBC survival from the aspect of miRNA regulation.

## RESULTS

### Genes with Informative APA Events are associated with Breast Cancer

The random sampling and univariate COX regression analysis identified 1,789 informative APA events (Figure 1A) including 881 events in breast cancer related genes as shown in Figure 1B. These events were enriched in ten pathways associated with breast cancer, APA, and miRNA regulation (Figure 1C). For example, rRNA processing can be promoted for tumor formation in breast cancer cells ([34]). Nucleic acid mediated communication between cancer cells and stroma partially promote the emergence, metastasis, and resistance of breast cancer ([35]). Nucleocytoplasmic transport may affect estrogen receptor alpha (ERα) to induce the expression of genes associated with the expansion of breast cancer ([36]). Some mechanisms of chromatin remodeling such as the methylation of *H3K27*, *H3K9*, and *H3K4* were demonstrated to be associated with breast cancer in previous studies ([37]). Moreover, the pathways that enriched miRNAs and APA ([13, 38]) guided us to study the mechanisms of APA events related to TNBC survival from the aspect of miRNA regulation in the following part.

**FIGURE 1.**
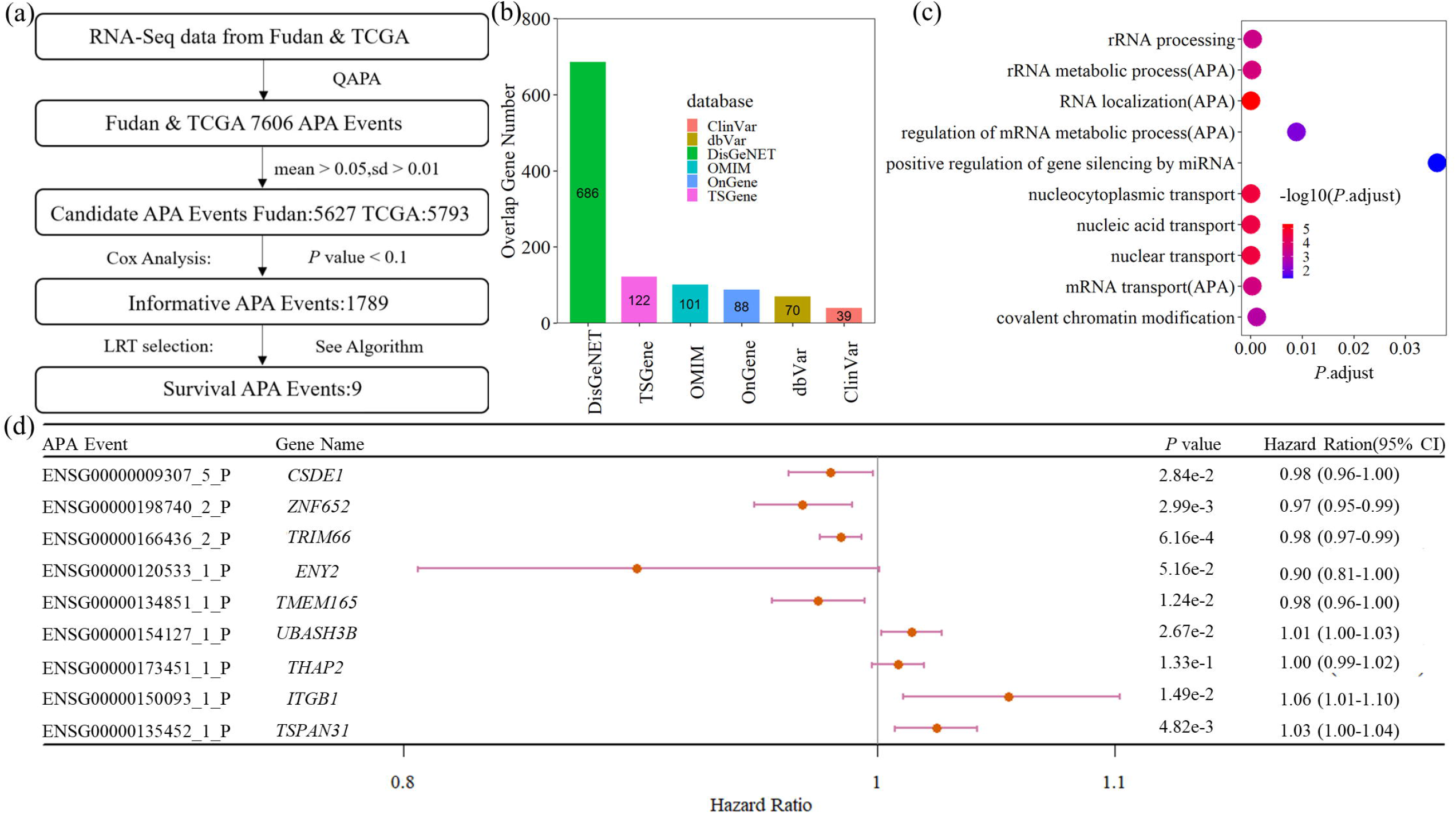
Screening process and disease association analyses of informative and survival APA events. (a) Flow chart shows the screening pipeline. (b) Histogram displays the overlap between informative APA events and breast cancer related genes from popular tumor databases. (c) Bubble charts present breast cancer and APA related enrichment pathways of genes with informative APA events. (d) Forest plot illustrates Cox Multivariate Regression Analysis results of survival APA events in merged datasets.

### Nine APAs are Potential Factors for TNBC Survival Prognosis

The forward-selected LRT identified nine survival APA events. They either presented unfavorable (hazard ratio (HR) > 1) or favorable (HR < 1) effects on TNBC survival prognosis (Figure 1D). To further understand the roles of these APA events in TNBC prognosis, we constructed a risk signature with Fudan dataset as shown in Figure 2A. Based on this signature, we determined 0.7284 as the optimal threshold between the high and low risk groups (Figure 2B). The performance of this risk signature was assessed by KM survival curves in Fudan and TCGA datasets. The results in Figure 2C–D, Figure 3, and Extended Data Figure2 showed that this signature could significantly classify the patients with high or low survival risks in almost each clinical category.

**FIGURE 2.**
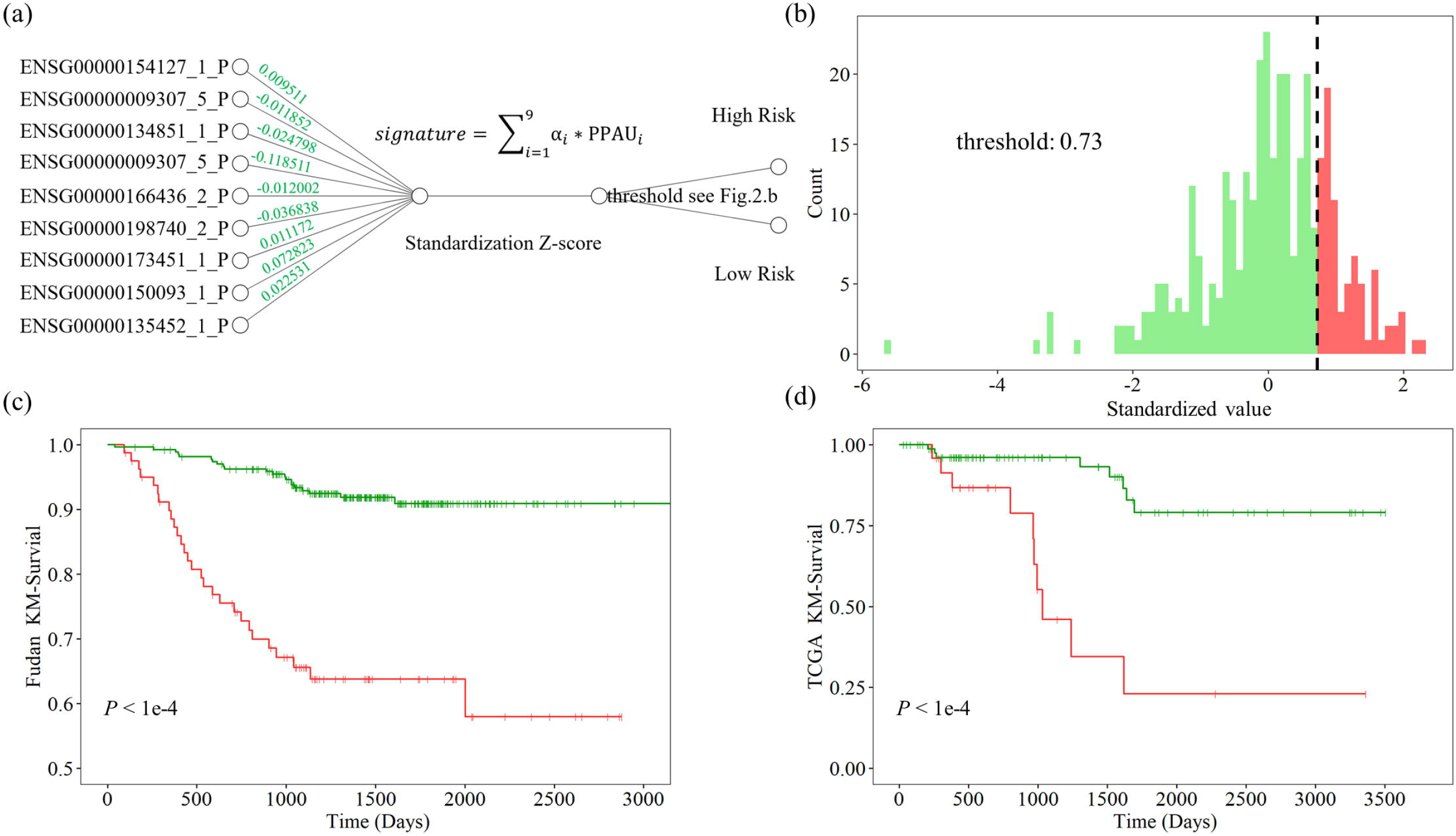
The calculation method and assessment of the risk signature. (a) Network diagram shows the calculation process of a risk signature. The risk signature is obtained by normalized weighted sum of survival APAs’ PPAU. Patients were then divided into high and low risk groups based on threshold. (b) Frequency histogram displays the optimal threshold and proportion of two groups in Fudan dataset. (c-d) Kaplan Meier survival curves of the risk signatures based on the samples from Fudan and TCGA datasets.

**FIGURE 3.**
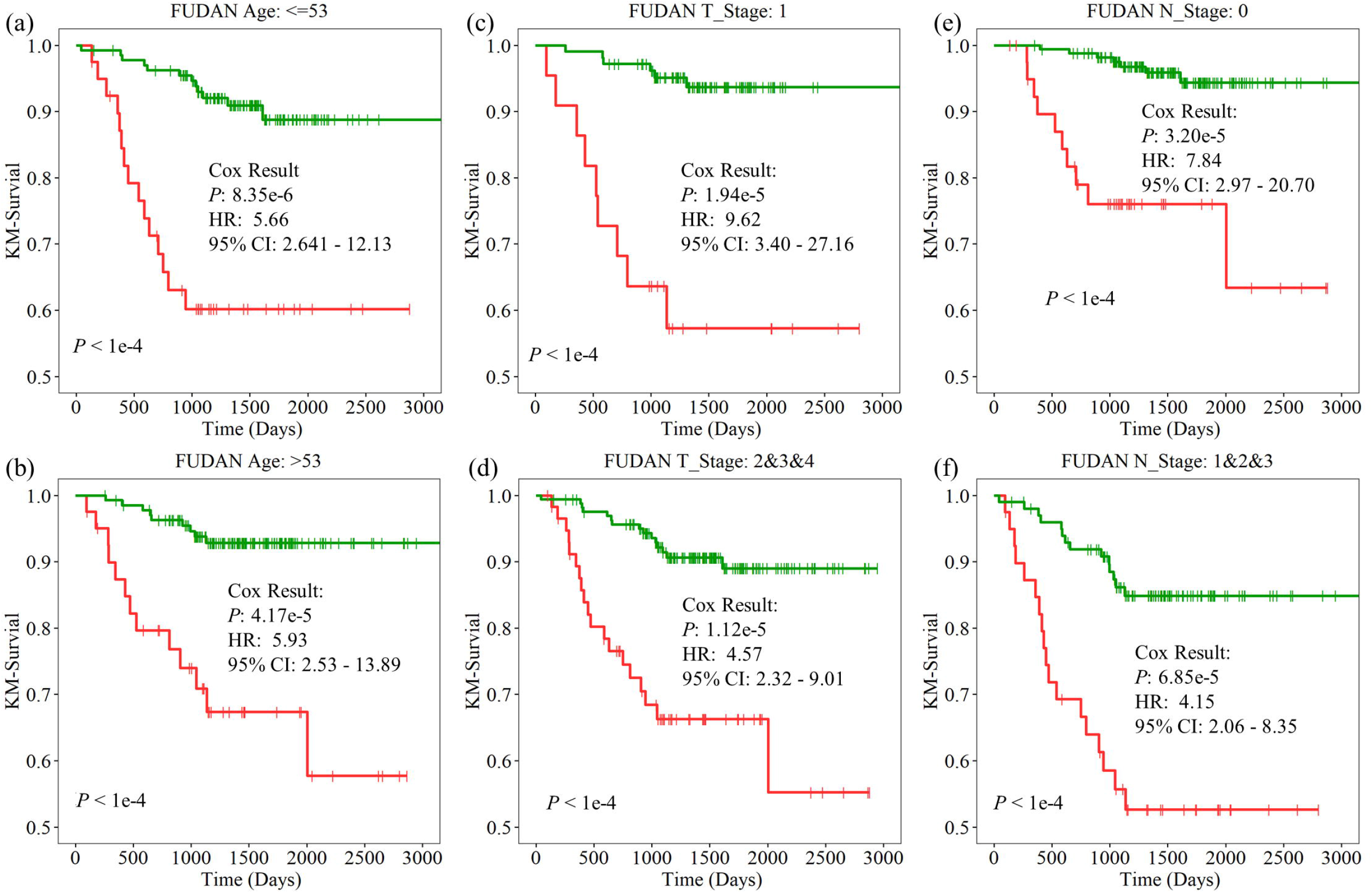
The usefulness of the APA risk signature in RFS prognosis under different clinical groups. Kaplan Meier survival curves in the groups of (a) Age < = 53, (b) Age > 53, (c) T1, (d)T2-4, (e) N0, and (f) N1-3 of Fudan dataset. It shows that APA risk signature still had the power to significantly classify patients into the high and low risk groups even under different clinical conditions. The results of TCGA database are displayed in Extended Data FIGURE 2.

In addition, we combined this risk signature and clinical information for the prediction of TNBC prognosis. As shown in Figure 4, the model with APA risk signature alone performed better than that used clinical information alone. Moreover, the combined features had better prognosis ability, realizing AUC values of 0.84 (0.78-0.90), 0.81 (0.73-0.89), 0.93 (0.86-1.00), and 0.86 (0.72-1.00) for the prognosis of 3-year RFS in Fudan dataset, 5-year RFS in Fudan dataset, 3-year RFS in TCGA dataset, 5-year RFS in TCGA dataset.

**FIGURE 4.**
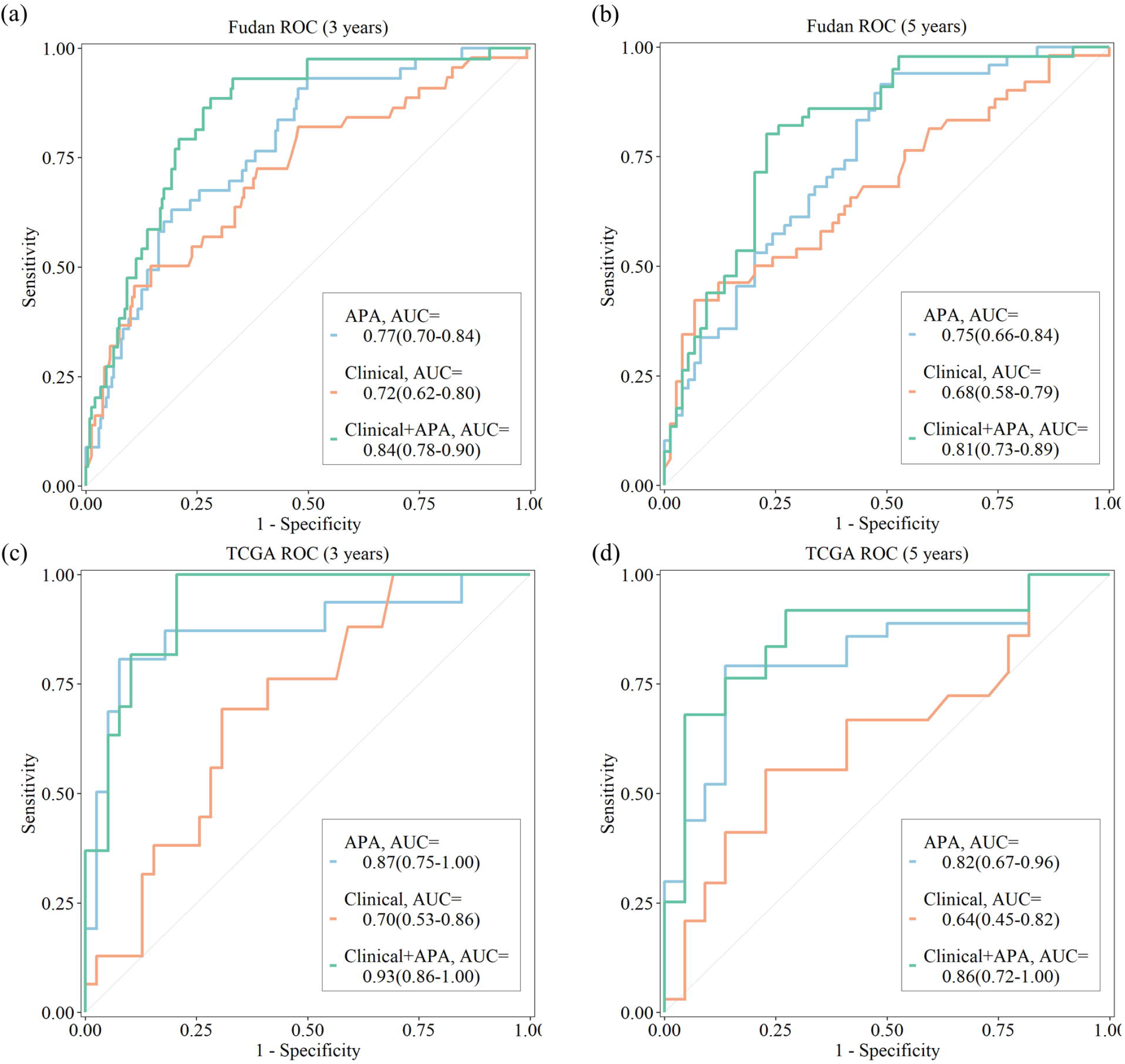
The performance of APA risk signature combined with clinical information. The performances of clinical information, APA risk signature, and the combined features are shown (a) for 3-year RFS in Fudan dataset, (b) for 5-year RFS in Fudan dataset, (c) for 3-year RFS in TCGA dataset, and (d) for 5-year RFS in TCGA dataset. It reveals that APA risk signature is more powerful in predicting relapse status.

All the above analyses demonstrated the potential of the nine survival APA events in TNBC survival prognosis. It revealed the necessity to study the possible regulatory pathways of these survival APA events further.

### APA Affects TNBC Prognosis by Altering the miRNA Regulation on its Host Gene

Among these nine survival APA events, four can affect the expression of their host genes by leading to the loss of miRNA regulation. The hypothesis was supported by the detection of miRNA targets using TargetScan and the correlation analysis of PPAUs with expression levels in both datasets of TCGA and Fudan (Figure 5A-B and Extended Data Figure 3B-H).

**FIGURE 5.**
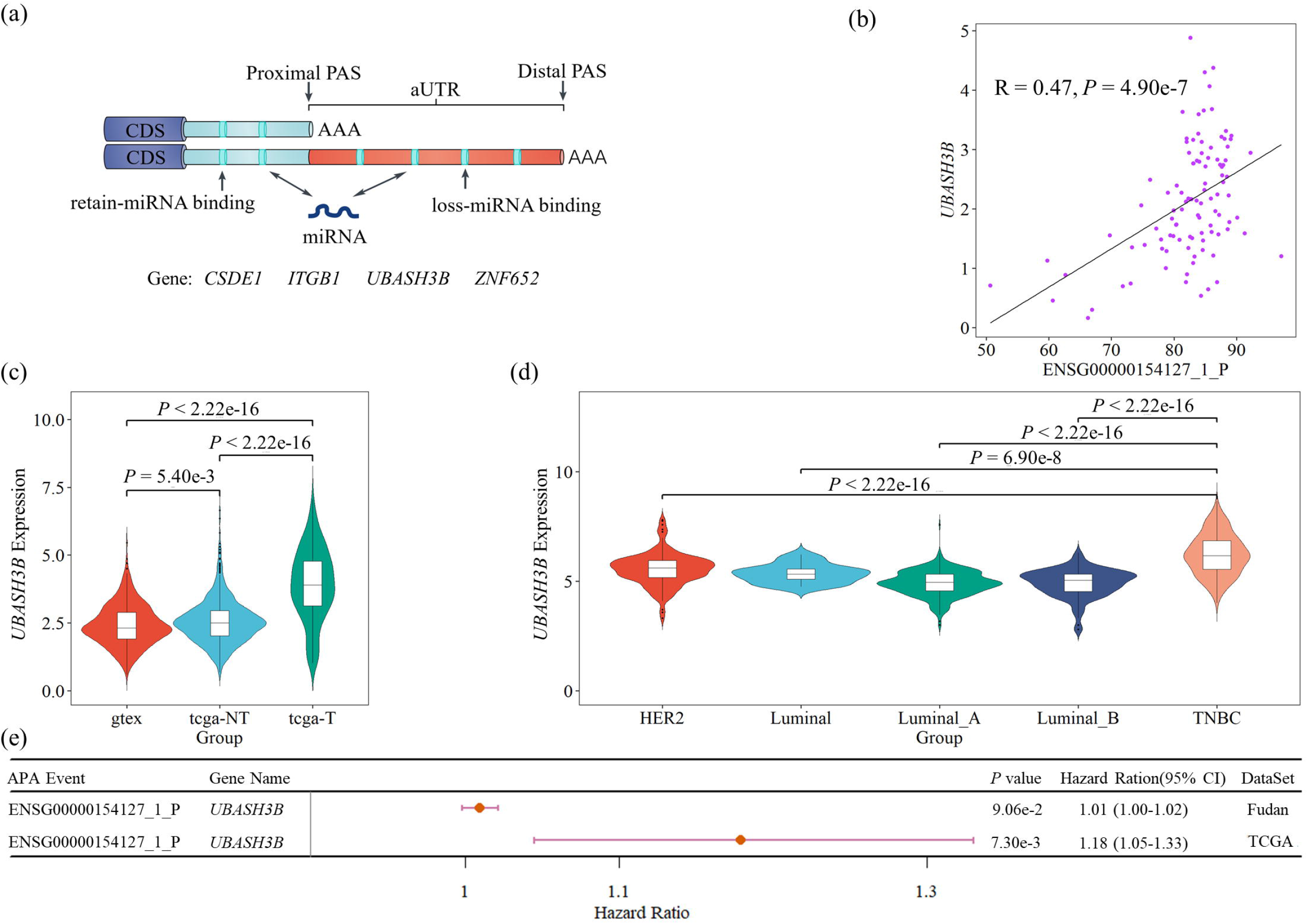
APA event on *UBASH3B* may affect the expression level of its host gene. (a) APA leads to the loss of miRNA bindings. (b) The relationship between the expression of *UBASH3B* and the PPAU of its proximal APA event in TCGA dataset. (c) Violin plot reveals the expression of *UBASH3B* was gradually and significantly increased in breast cancer (tcga-NT), and TNBC tissues (tcga-T). (d) Violin plot shows that the expression level of *UBASH3B* in TNBC is higher than other breast cancer subtypes from GENT2 database. (e) Forest plot shows that the hazard ratio of the APA event in *UBASH3B* is greater than one in two datasets. Other associations can be seen in Extended Data FIGURE 3.

Among these four events, APA event in *UBASH3B* showed the highest association with its host gene (R = 0.47). Thus, it was selected to study further for its potential mechanisms in TNBC survival. From previous studies, we got the knowledge that *UBASH3B* was overexpressed in TNBC to promote the invasion and metastasis of tumors ([39]). Our analysis results also supported this discovery with higher expression in more aggressive breast cancer samples (Figure 5C), the highest expression in TNBC among all the breast cancer subtypes (Figure 5D), and the unfavorable effects (HR>1) of this APA event on TNBC survival in both datasets (Figure 5E). The above results suggested that this APA event caused the increased expression of *UBASH3B* by altering the miRNA regulation to probably increase the risk of TNBC patients.

### APA Altered the Competition between *UBASH3B* and *ZNF652* through Interfering in miRNA Regulation

Since APA events can affect the expression of their host genes, then other genes competing with their host genes may also be altered by these APA events. Thus, for the survival APA event in *UBASH3B*, we next explored its effects on other genes competed with its host gene. After the analysis, *ZNF652* was identified as a competing gene of *UBASH3B* (Extended Data Figure 4A & Figure 6A). Their competition relationships were supported by their negative expression associations in both two datasets. Then the APA event in *UBASH3B* may indirectly affect *ZNF652*, which was backed up with the remarkable relationships between the expression levels of *ZNF652* and the PPAUs of *UBASH3B* in TCGA dataset (Figure 6B). All these results indicated that the APA-mediated miRNA regulation not only enhanced the expression of *UBASH3B*, but also competitively inhibited the expression of *ZNF652*.

**FIGURE 6.**
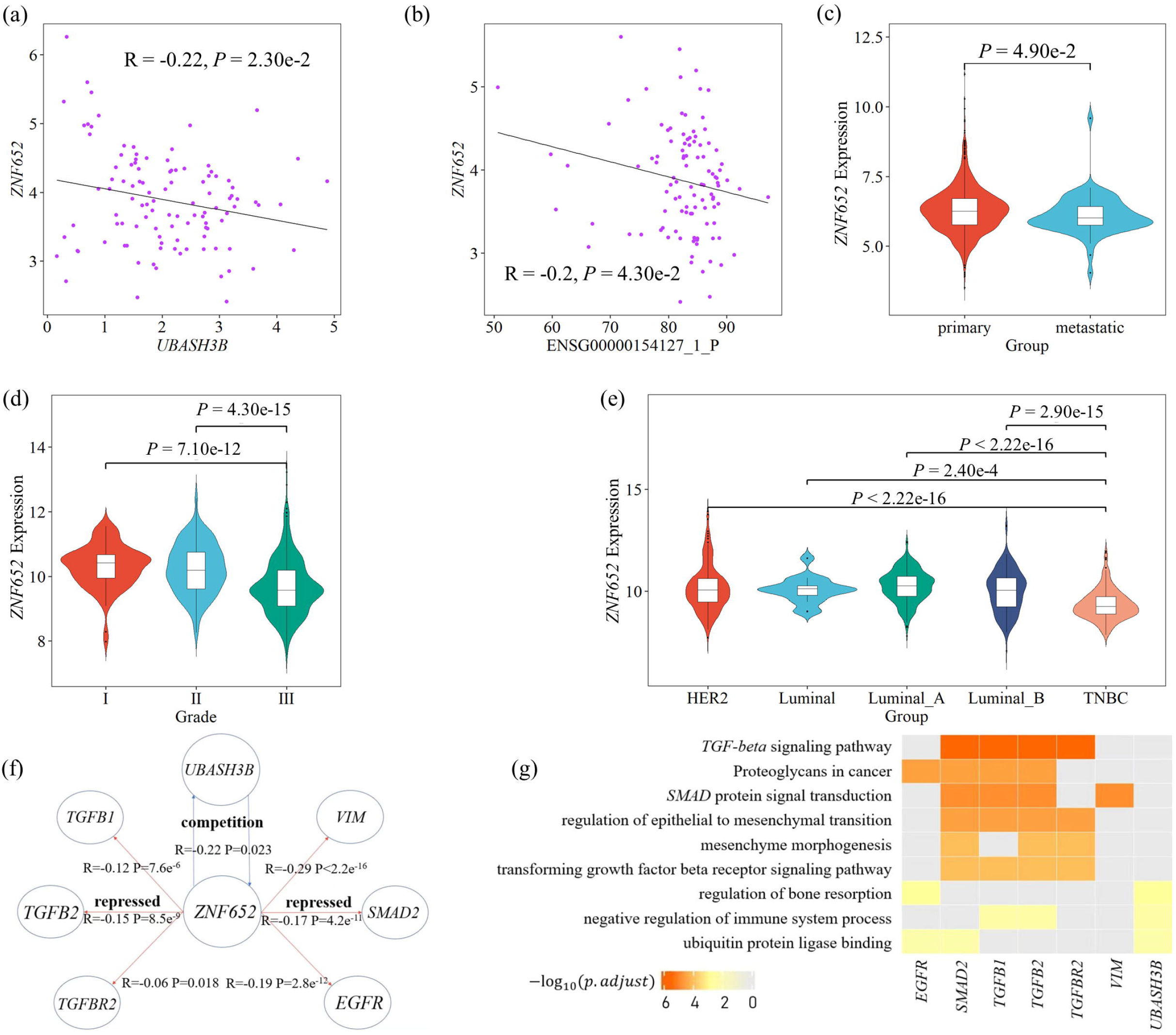
Downstream analysis of the APA event in *UBASH3B*. (a) The relationship of gene expression level between *ZNF652* and *UBASH3B* in TCGA dataset. (b) The relationship between the expression level of *ZNF652* and APA event in *UBASH3B*. (c) Violin plot shows the significantly lower expressions of *ZNF652* in TCGA metastatic tissue than in primary tissue. (d) Violin plot shows that the expression level of *ZNF652* decreases with increasing disease stage from GENT2 breast cancer dataset. (e) Violin plot shows that the expression level of *ZNF652* in TNBC is higher than other breast cancer subtypes from GENT2 breast cancer dataset. (f) Breast cancer-related gene expression regulatory network associated with the APA event in *UBASH3B*. *ZNF652* affected by the APA event in *UBASH3B* directly repressed some key driver genes of invasion and metastasis such as *TGFB1*, *TGFB2*, *TGFBR2*, *VIM*, *SMAD2*, and *EGFR.* (f) GO and KEGG enrichment results of the genes in the network were related to breast cancer. Other associations can be seen in Extended Data FIGURE 4.

From previous research, *ZNF652* is a direct regulator of epithelial–mesenchymal transition (EMT) and represses six key drivers of invasion and metastasis, including *TGFB1*, *TGFB2*, *TGFBR2*, *EGFR*, *SMAD2*, and *VIM* ([40]). During our analyses, we also discovered the role of *ZNF652* to degrade tumor metastasis and cancer progression, since the expression levels of *ZNF652* were lower in metastatic samples, severe samples, and TNBC subtype (Figure 6C-E), and was negatively correlated with the six key drivers (Figure 6F, Extended Data Figure 4C-H). Further, the enrichment analysis of *UBASH3B*, *ZNF652*, and the six key drivers were also involved in nine breast cancer-related pathways (Figure 6G). The above results displayed that this APA may alter the miRNA competition relationships between *UBASH3B* and *ZNF652* to further attenuate the inhibition of *ZNF652* on key drivers of invasion and metastasis.

## DISCUSSION

TNBC has the highest rates of recurrence, metastasis, and mortality, and limited effective targeted therapy. There are studies which proposed several TNBC related biomarkers involving in gene expression ([41, 42]), signal transduction pathway ([43]), DNA damage response ([44–46]), tumor microenvironment ([47]), and miRNA regulation ([48]). Although these biomarkers have shown certain effects in some studies, we still need to research on potential biomarkers from more aspects due to the strong heterogeneity of TNBC. In this study, we identified nine survival APA events. Their risk signature performed well (Fudan: *P*_KM_ = 6.62e-12; TCGA: *P*_KM_ = 5.38e-6) in samples with different races. The model composed of these nine survival APA events had better predictive power than common clinical indicators and previous signature (The area under ROC curves of the signatures at 5 years for RFS in the TCGA: 0.82 vs 0.73) ([49]). Finally, the APA event in *UBASH3B* was recognized as a new potential biomarker which disturbed miRNA regulation on its host gene and other competing tumor genes to play its roles in TNBC.

Before the identification of survival APA events, we used QAPA to identify APA events. It included more than half of the events detected by another tool (DaPars), as shown in Extended Data Figure 5A. Moreover, the mean APA characteristic values estimated by these two tools showed strong correlations (R = −0.73, *P* = 2.2e-16) in Extended Data Figure 5B. All these two analyses demonstrated the reliability of our pipeline for APA identification.

Further, we found the effect of APA event in *UBASH3B* on the expression of its host gene, through altering miRNA regulation. In the breast cancer cell lines of Cancer Cell Line Encyclopedia (CCLE), we also found that this APA event was significantly correlated with the expression of *UBASH3B* (P < 0.05, Extended Data Figure 6A). It revealed the correctness and applicability of this biomarker.

Overall, this study focused on the impact of APA-mediated miRNA regulation on TNBC prognosis. However, we could not ignore that there are other possible mechanisms such as RNA-binding proteins (RBPs) regulation ([13]), and single nucleotide polymorphisms (SNPs) ([50]) possibly involved in the APA functions. The impacts of RBPs on APA functions mainly include two aspects. On one hand, APAs of RBP targets may lead to the alterations of RBP binding regions to affect the expression of their host genes. For example, the APA event in *TRIM66* may cause the loss of 20 RBP targets on its host gene to affect its expression in Extended Data Figure 7A-B. On the other hand, APA events on RBPs may affect the regulation on their targeted genes. As shown in Extended Data Figure 7C-D, APA events in two RNA binding proteins may affect the expression of their targets, such as *UBASH3B*. The effect of APA-mediated RBP regulation on the prognosis of TNBC will be our next plan.

Further, we also discovered that APA events could be possibly affected by SNPs. In the CAFuncAPA (https://relab.xidian.edu.cn/CAFuncAPA/) database, for the APA event on *UBASH3B* discussed in this study, we discovered its differential Percentage of Distal polyA site Usage Index (PDUI) values (ANOVA: *P* = 0.0111) among the three genotyping groups. This analysis about the effect of SNPs on APA will be our second research plan.

## Supporting information

Supplementary Algorithm and Figure

## AUTHOR CONTRIBUTIONS

SW, CH conceived the project. XW and CH contributed to the acquisition of data. CH developed the algorithm. CH performed the analysis and wrote the manuscript. SW, CH, LH, and XZ contributed to the critical revision of the manuscript, SW and LH contributed the funding acquisition, LH and XZ contributed to the supervision of this project. All authors contributed to the article and approved the submitted version.

## ACKNOWLEDGMENTS

We thank the members of the School of Life Sciences and Technology for valuable discussions.

## REFERENCES

1. Foulkes WD, Smith IE, Reis-Filho JS. Triple-negative breast cancer, New England Journal of Medicine 2010;363:1938–1948.

2. Cleator S, Heller W, Coombes RC. Triple-negative breast cancer: therapeutic options, The lancet oncology 2007;8:235–244.

3. Irvin Jr WJ, Carey LA. What is triple-negative breast cancer?, European journal of cancer 2008;44:2799–2805.

4. Synnott N, Murray A, McGowan P et al. Mutant p53: a novel target for the treatment of patients with triple□negative breast cancer?, International journal of cancer 2017;140:234–246.

5. Saloustros E, Nikolaou M, Kalbakis K et al. Weekly paclitaxel and carboplatin plus bevacizumab as first-line treatment of metastatic triple-negative breast cancer. A multicenter phase II trial by the Hellenic oncology research group, Clinical Breast Cancer 2018;18:88–94.

6. Becattini C, Agnelli G, Schenone A et al. Aspirin for preventing the recurrence of venous thromboembolism, New England journal of medicine 2012;366:1959–1967.

7. Miller K, Wang M, Gralow J et al. Paclitaxel plus bevacizumab versus paclitaxel alone for metastatic breast cancer, New England journal of medicine 2007;357:2666–2676.

8. Niemeier LA, Dabbs DJ, Beriwal S et al. Androgen receptor in breast cancer: expression in estrogen receptor-positive tumors and in estrogen receptor-negative tumors with apocrine differentiation, Modern Pathology 2010;23:205–212.

9. He J, Peng R, Yuan Z et al. Prognostic value of androgen receptor expression in operable triple-negative breast cancer: a retrospective analysis based on a tissue microarray, Medical oncology 2012;29:406–410.

10. Park S, Koo J, Park H et al. Expression of androgen receptors in primary breast cancer, Annals of oncology 2010;21:488–492.

11. Gucalp A, Traina TA. Targeting the androgen receptor in triple-negative breast cancer, Current problems in cancer 2016;40:141–150.

12. da Silva JL, Nunes NCC, Izetti P et al. Triple negative breast cancer: A thorough review of biomarkers, Critical reviews in oncology/hematology 2020;145:102855.

13. Tian B, Manley JL. Alternative polyadenylation of mRNA precursors, Nature reviews Molecular cell biology 2017;18:18–30.

14. Komini C, Theohari I, Lambrianidou A et al. PAPOLA contributes to cyclin D1 mRNA alternative polyadenylation and promotes breast cancer cell proliferation, Journal of Cell Science 2021;134:jcs252304.

15. Tamaddon M, Shokri G, Hosseini Rad SMA et al. Involved microRNAs in alternative polyadenylation intervene in breast cancer via regulation of cleavage factor “CFIm25”, Scientific reports 2020;10:1–11.

16. Jiang Y-Z, Ma D, Suo C et al. Genomic and transcriptomic landscape of triple-negative breast cancers: subtypes and treatment strategies, Cancer Cell 2019;35:428–440. e425.

17. Ha KC, Blencowe BJ, Morris Q. QAPA: a new method for the systematic analysis of alternative polyadenylation from RNA-seq data, Genome biology 2018;19:1–18.

18. Patro R, Duggal G, Love MI et al. Salmon provides fast and bias-aware quantification of transcript expression, Nature methods 2017;14:417–419.

19. Dobin A, Davis CA, Schlesinger F et al. STAR: ultrafast universal RNA-seq aligner, Bioinformatics 2013;29:15–21.

20. Li B, Dewey CN. RSEM: accurate transcript quantification from RNA-Seq data with or without a reference genome, BMC Bioinformatics 2011;12:1–16.

21. Yu G, Wang L-G, Han Y et al. clusterProfiler: an R package for comparing biological themes among gene clusters, Omics: a journal of integrative biology 2012;16:284–287.

22. Piñero J, Ramírez-Anguita JM, Saüch-Pitarch J et al. The DisGeNET knowledge platform for disease genomics: 2019 update, Nucleic acids research 2020;48:D845–D855.

23. Liu Y, Sun J, Zhao M. ONGene: a literature-based database for human oncogenes, Journal of Genetics and Genomics 2017;44:119–121.

24. Amberger JS, Bocchini CA, Schiettecatte F et al. OMIM. org: Online Mendelian Inheritance in Man (OMIM®), an online catalog of human genes and genetic disorders, Nucleic acids research 2015;43:D789–D798.

25. Zhao M, Kim P, Mitra R et al. TSGene 2.0: an updated literature-based knowledgebase for tumor suppressor genes, Nucleic acids research 2016;44:D1023–D1031.

26. Lappalainen I, Lopez J, Skipper L et al. DbVar and DGVa: public archives for genomic structural variation, Nucleic acids research 2012;41:D936–D941.

27. Landrum MJ, Lee JM, Benson M et al. ClinVar: public archive of interpretations of clinically relevant variants, Nucleic Acids Research 2016;44:D862–D868.

28. Harrell Jr FE, Harrell Jr MFE, Hmisc D. Package ‘rms’, Vanderbilt University 2017;229.

29. Wang L, Hu X, Wang P et al. The 3’ UTR signature defines a highly metastatic subgroup of triple-negative breast cancer, Oncotarget 2016;7:59834.

30. Camp RL, Dolled-Filhart M, Rimm DL. X-tile: a new bio-informatics tool for biomarker assessment and outcome-based cut-point optimization, Clinical cancer research 2004;10:7252–7259.

31. Sandberg R, Neilson JR, Sarma A et al. Proliferating cells express mRNAs with shortened 3’untranslated regions and fewer microRNA target sites, Science 2008;320:1643–1647.

32. Mayr C, Bartel DP. Widespread shortening of 3’ UTRs by alternative cleavage and polyadenylation activates oncogenes in cancer cells, Cell 2009;138:673–684.

33. Lewis BP, Burge CB, Bartel DP. Conserved seed pairing, often flanked by adenosines, indicates that thousands of human genes are microRNA targets, Cell 2005;120:15–20.

34. Zhang Y, Baysac KC, Yee L-F et al. Elevated DDX21 regulates c-Jun activity and rRNA processing in human breast cancers, Breast Cancer Research 2014;16:1–18.

35. Yu Dd, Wu Y, Shen Hy et al. Exosomes in development, metastasis and drug resistance of breast cancer, Cancer science 2015;106:959–964.

36. Tecalco-Cruz AC, Pérez-Alvarado IA, Ramírez-Jarquín JO et al. Nucleo-cytoplasmic transport of estrogen receptor alpha in breast cancer cells, Cellular Signalling 2017;34:121–132.

37. Wang GG, Allis CD, Chi P. Chromatin remodeling and cancer, Part I: Covalent histone modifications, Trends in molecular medicine 2007;13:363–372.

38. Goering R, Engel KL, Gillen AE et al. LABRAT reveals association of alternative polyadenylation with transcript localization, RNA binding protein expression, transcription speed, and cancer survival, BMC genomics 2021;22:1–27.

39. Lee ST, Feng M, Wei Y et al. Protein tyrosine phosphatase UBASH3B is overexpressed in triple-negative breast cancer and promotes invasion and metastasis, Proceedings of the National Academy of Sciences 2013;110:11121–11126.

40. Neilsen PM, Noll JE, Mattiske S et al. Mutant p53 drives invasion in breast tumors through up-regulation of miR-155, Oncogene 2013;32:2992–3000.

41. Viale G, Group tIBCS, Regan MM et al. Predictive value of tumor Ki-67 expression in two randomized trials of adjuvant chemoendocrine therapy for node-negative breast cancer, JNCI: Journal of the National Cancer Institute 2008;100:207–212.

42. Blows FM, Driver KE, Schmidt MK et al. Subtyping of breast cancer by immunohistochemistry to investigate a relationship between subtype and short and long term survival: a collaborative analysis of data for 10,159 cases from 12 studies, PLoS medicine 2010;7:e1000279.

43. Liu D, He J, Yuan Z et al. EGFR expression correlates with decreased disease-free survival in triple-negative breast cancer: a retrospective analysis based on a tissue microarray, Medical oncology 2012;29:401–405.

44. Couch FJ, Hart SN, Sharma P et al. Inherited mutations in 17 breast cancer susceptibility genes among a large triple-negative breast cancer cohort unselected for family history of breast cancer, Journal of clinical oncology 2015;33:304.

45. Sharma P, Klemp JR, Kimler BF et al. Germline BRCA mutation evaluation in a prospective triple-negative breast cancer registry: implications for hereditary breast and/or ovarian cancer syndrome testing, Breast cancer research and treatment 2014;145:707–714.

46. Telli ML, Timms KM, Reid J et al. Homologous Recombination Deficiency (HRD) Score Predicts Response to Platinum-Containing Neoadjuvant Chemotherapy in Patients with Triple-Negative Breast CancerHRD Predicts Response to Platinum Therapy in TNBC, Clinical cancer research 2016;22:3764–3773.

47. Dieci M, Criscitiello C, Goubar A et al. Prognostic value of tumor-infiltrating lymphocytes on residual disease after primary chemotherapy for triple-negative breast cancer: a retrospective multicenter study, Annals of oncology 2014;25:611–618.

48. Lü L, Mao X, Shi P et al. MicroRNAs in the prognosis of triple-negative breast cancer: A systematic review and meta-analysis, Medicine 2017;96.

49. Zhang Y, Wang Y, Li C et al. Systemic analysis of the prognosis-associated alternative polyadenylation events in breast cancer, Frontiers in Genetics 2020;11:590770.

50. Thomas LF, Sætrom P. Single nucleotide polymorphisms can create alternative polyadenylation signals and affect gene expression through loss of microRNA-regulation 2012.

