## Supplementary Algorithm and Figure for "Alternative Polyadenylation alters UBASH3B-ZNF652 competition to involve in Triple-Negative Breast Cancer"

**SUPPLEMENTARY MATERIAL**

**1. Algorithm**

**Algorithm: Informative and Survival APA Events Screening Process**

**Input:** Two Dataset PPAU Arrays (set1, set2), echo = 1000, max_l = 0.05

**Output:** APA Screening Result (Informative_APA_sites)

1: create a list (dim = echo);

2: **for** i = 1; i ≤ echo; i ++ **do**

3: subset1 ← set1 (randomly drawing 70% from all set1's samples);

4: temp_result1 ← subset1 (Univariate Cox Regression Analysis);

5: subset2 ← set2 (randomly drawing 70% from all set2's samples);

6: temp_result2 ← subset2 (Univariate Cox Regression Analysis);

7: APA_sites1 ← temp_result1 (filtering with *P* ≤ 0.1);

8: APA_sites2 ← temp_result2 (filtering with *P* ≤ 0.1);

9: list[i] = APA_sites1 ∩ APA_sites2;

10: **end for**

11: Count the number of occurrences of each APA site from list (the max number ≤ echo);

12: sort_result ← Sort APA sites according to the number of occurrences;

13: set ← set1 ∪ set2;

14: Add sort_result[1] in Informative_APA_sites;

15: Likelihood Ratio Test by forward selection variables:

16: **for** j = 2; j ≤ max_l*size(sort_result); j ++ **do**

17:	model_current ← cph (Informative_APA_sites)

18:	model_new ← cph (Informative_APA_sites ∪ sort_result[j])

19: lr_result ← lr_test (model_current, model_new)

20: if lr_result$*P*.value ≤ 0.05 then

21: Add sort_result[j] in Informative_APA_sites;

22: **end if**

23: **end for**

24: < cph: Cox Proportional Hazards Mode >

25: < lr_test: Likelihood Ratio Test >

**2.** **Extended Data. FIGURE**

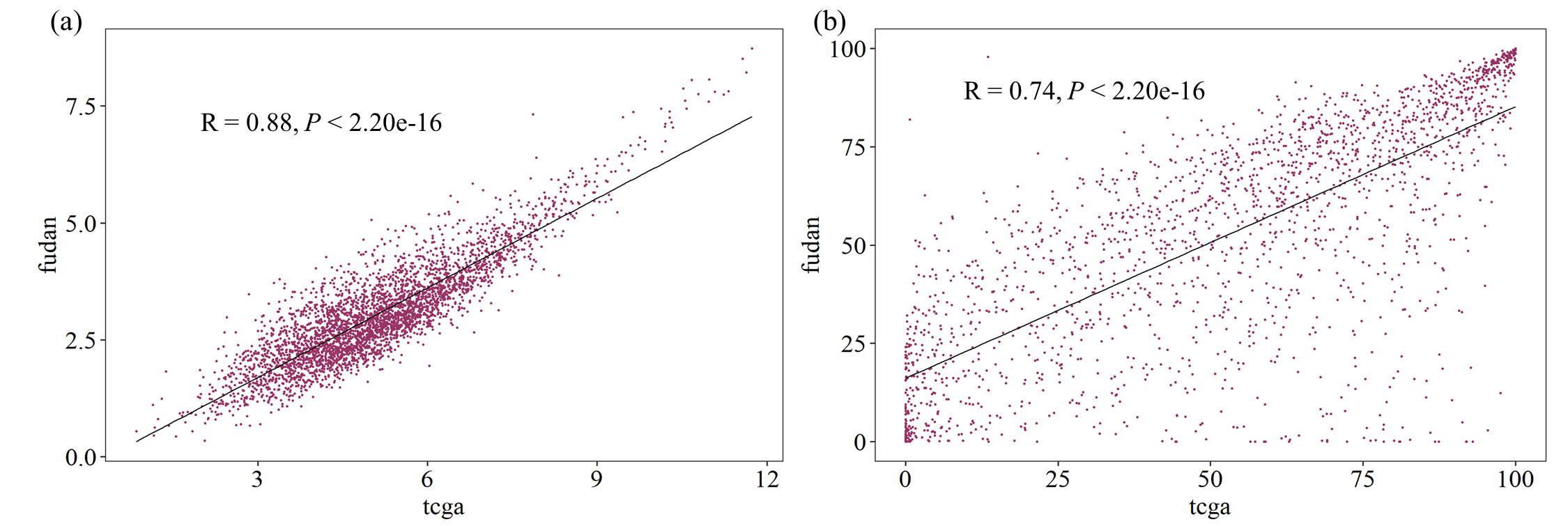

Extended Data. FIGURE 1 | The relationship of (a) the expression level of house-keeping genes, and (b) PPAU of proximal APA events’ between Fudan and TCGA dataset.

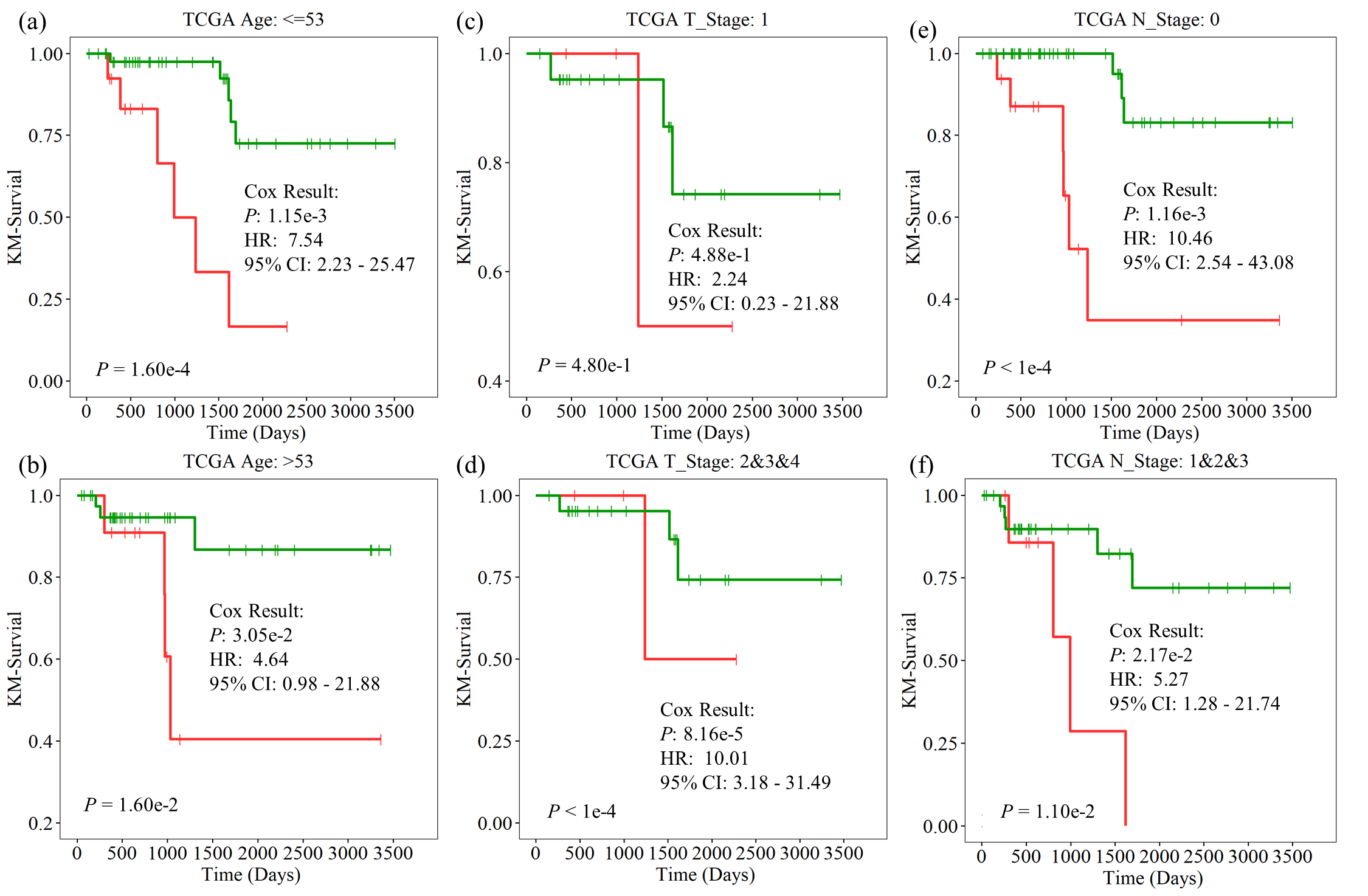

Extended Data. FIGURE 2 | The usefulness of the APA risk signature in RFS prognosis under different clinical groups. Kaplan Meier survival curves in the groups of (a) Age <= 53, (b) Age > 53, (c) T1, (d) T2-4, (e) N0, and (f) N1-3 of TCGA dataset. It shows that APA risk signature still had the power to significantly classify patients into the high and low risk groups even under different clinical conditions.

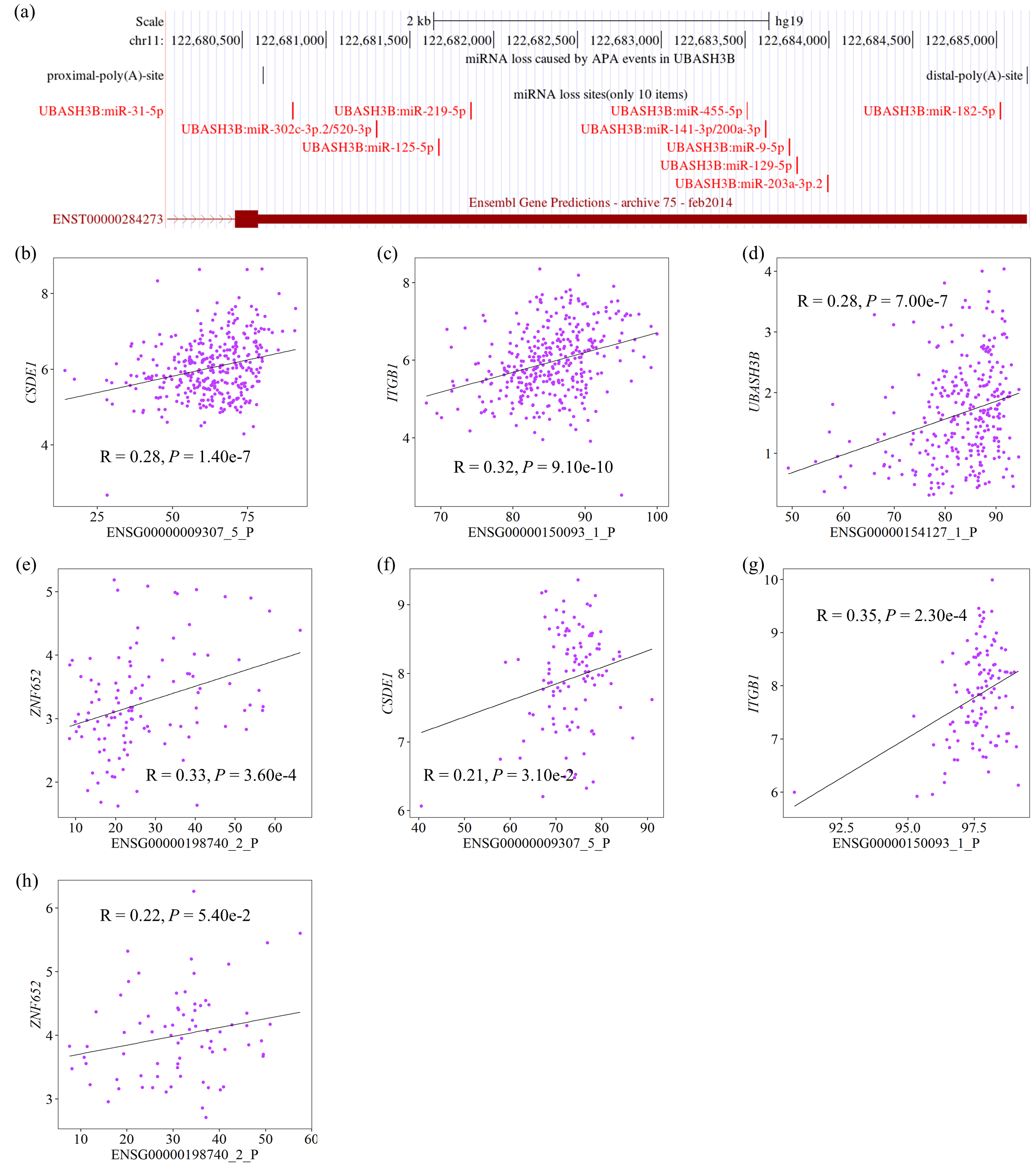

Extended Data. FIGURE 3 | APA events may regulate expression levels of their host genes. (a) The APA event in *UBASH3B* leads to the loss of miRNA bindings. The relationship of APA events in *CSDE1*, *ITGB1*, *UBASH3B*, and *ZNF652* with their host genes in (b-e) Fudan dataset and (f-h) TCGA dataset.

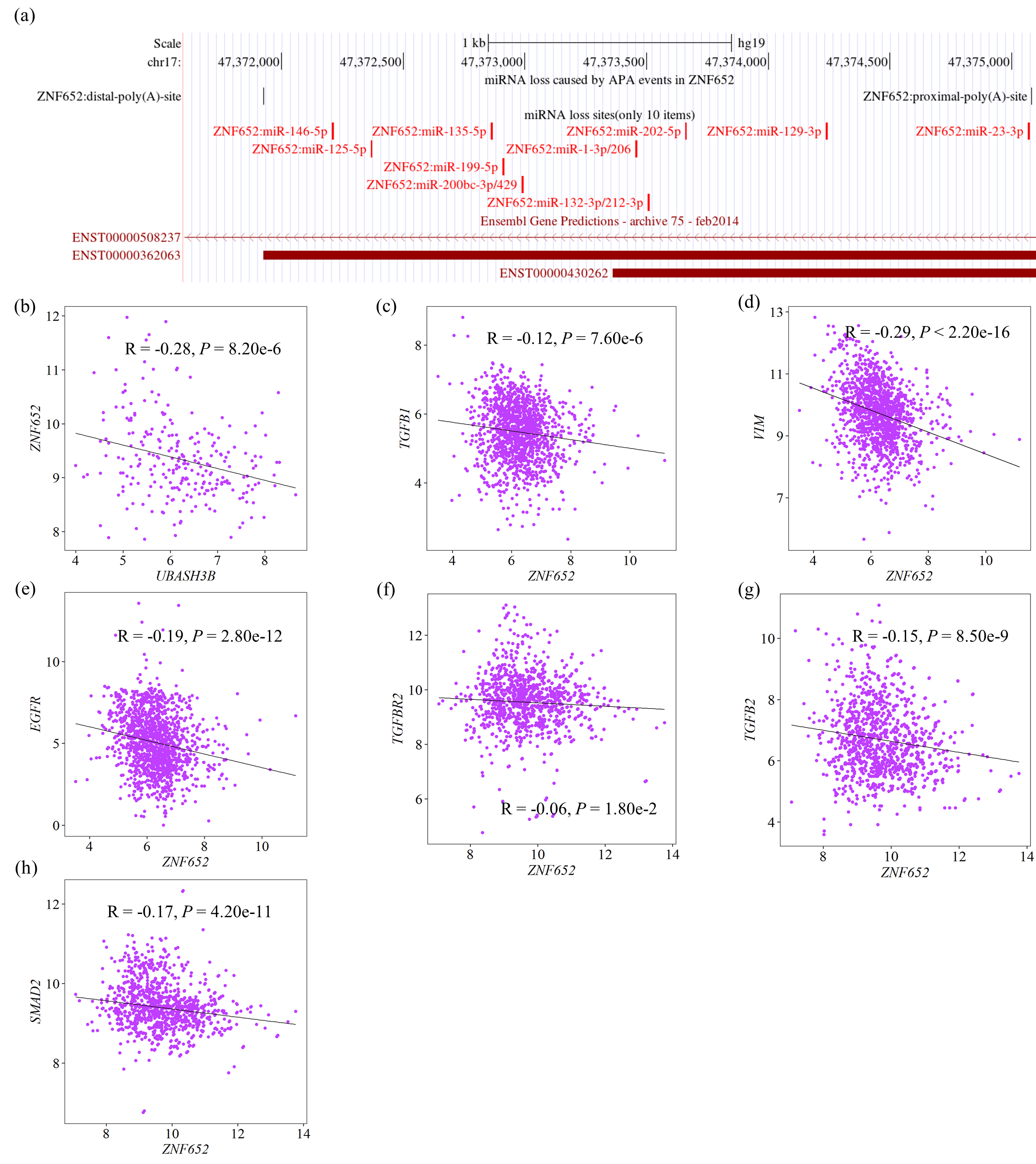

Extended Data. FIGURE 4 | *UBASH3B* inhibits the expression of *ZNF652* to attenuate its inhibition on key drivers of invasion and metastasis. (a) *ZNF652* competed with *UBASH3B* for miRNAs regulation. (b) The relationship between the expression level of *ZNF652* and *UBASH3B* in Fudan dataset. Correlations plots showed the relationship of the expression level between *ZNF652* and (c) *TGFB1*, (d) *VIM*, or (e) *EGFR* in TCGA primary breast cancer dataset. Correlations plots showed the relationship of the expression level between *ZNF652* and (f) *TGFBR2*, (g) *TGFB2*, or (h) *SMAD2* in GENT2 breast cancer dataset.

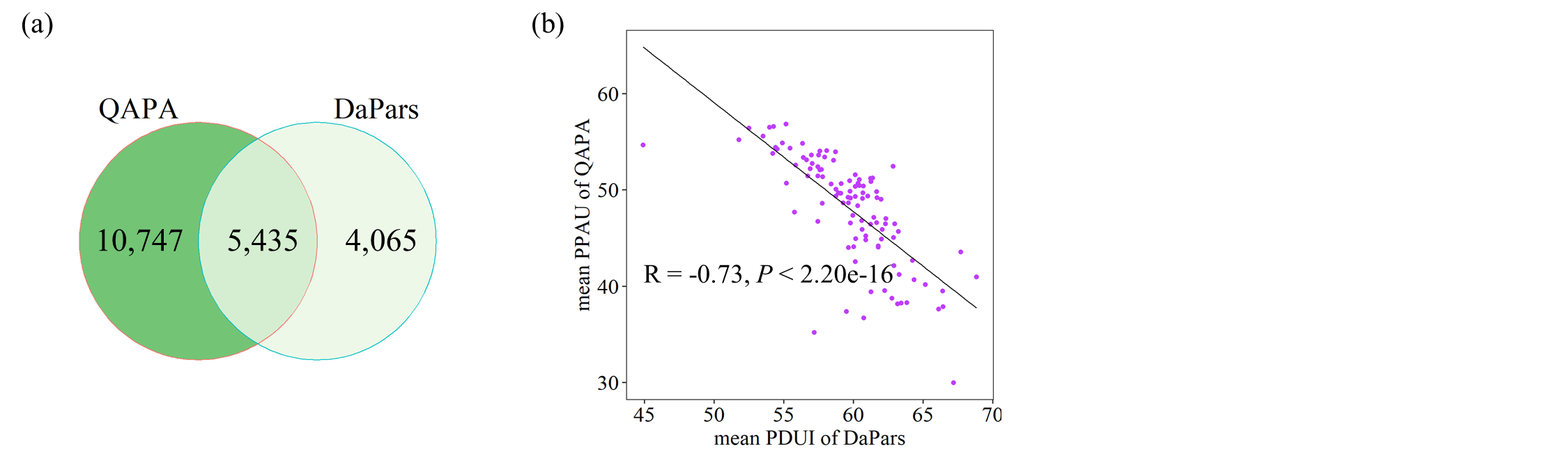

Extended Data. FIGURE 5 | The comparison of APA event identification pipeline in our study with DaPars used in The Cancer 3' UTR Atlas (TC3A). (a) Overlap of genes with the same APA site (with 50bp) identified by QAPA and DaPars. (b) Correlation of the mean values of APA (QAPA: PPAU DaPars: PDUI) identified by QAPA and DaPars.

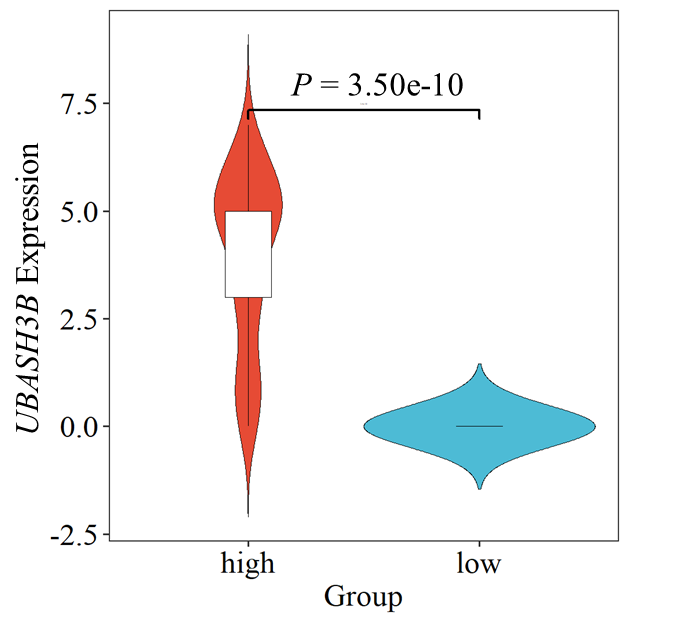

Extended Data. FIGURE 6 | Validation of biomarkers in CCLE dataset. The threshold of high and low grouping was determined by the mean value of PPAU for all samples.

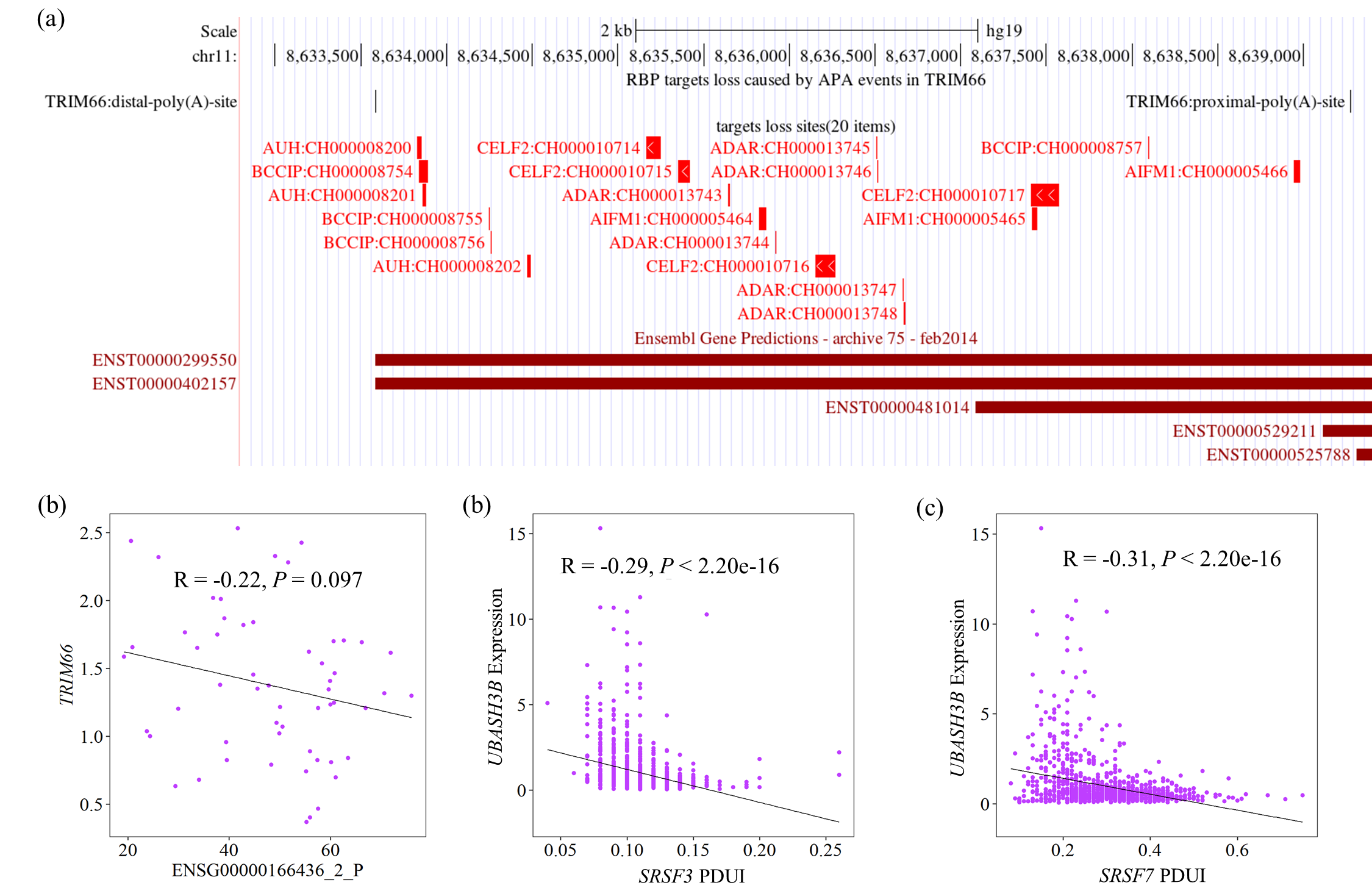

Extended Data. FIGURE 7 | The effect of APA-mediated RBP regulation on the prognosis of TNBC (a-b) APA event in *TRIM66* may cause the loss of RBP targets to affect the expression level of it in TCGA data. (c-d) It showed the correlations between the expression of *UBASH3B* and the PDUI of two RBPs, such as *SRSF3* and *SRSF7* in CAFuncAPA database.
